# Effects of urbanization on the microbial fungal communities from water, sediment, and amphibian hosts across a rural-to-urban waterway in Worcester, Massachusetts

**DOI:** 10.1101/2025.04.01.646624

**Authors:** Sara J. Wheeler, Manning DelCogliano, Brennan Hare, Alexander J. Bradshaw, Madison R. Hincher, Jasper P. Carleton, Philip J. Bergmann, Nathan Ahlgren, Javier F. Tabima

**Affiliations:** Biology Department, Clark University. Worcester, Massachusetts

**Keywords:** Urbanization, Fungal communities, Aquatic environments, Amphibian symbionts

## Abstract

In a world of increasingly urbanized environments, it is critical to understand the impact of urbanization on microbial communities, including fungal communities, as a measure of ecosystem health, and to document how these environments are changing. Aquatic environments, in particular, can be highly sensitive to urbanization with removal of local habitat and inputs of wastewater and contaminants, leading to displacement or extinction of natural flora, fauna, and funga. Aquatic microbial communities, especially fungal communities, are extremely complex and far less characterized compared to their macro counterparts. Here, we characterized the fungal microbiome, i.e. “mycobiome”, across multiple habitats— water, sediment, and frog fecal matter from three species (American bullfrog (*Lithobates catesbeianus*), Green frog (*Lithobates clamitans*), and Pickerel frog (*Lithobates palustris*)) — within the Tatnuck Brook waterway in Worcester, MA. The Tatnuck Brook waterway is a connected group of streams, ponds, and lakes that transition from protected areas at its headwaters, below the city’s drinking water reservoir, to highly urbanized regions within Worcester. Using metabarcode sequencing, we found that water, sediment, and frog gut habitats harbor distinct fungal communities, but all exhibit parallel shifts in diversity along an urbanization gradient. In particular, we identified fungal taxa from the amphibian mycobiome and environmental samples that exhibited a range of sensitivities to urbanization. These taxa, including *Basidiobolus*, *Cladosporium*, and *Lemonniera* in fecal and sediment samples, as well as *Candida*, which was found in all habitats, are potential indicators of shifts in aquatic ecosystems due to urbanization. Their responses to urban stressors may serve as a baseline for further studies of aquatic fungal communities.

**IMPORTANCE:** As cities grow, they can change nearby rivers, streams, and ponds in ways that affect plants, animals, and even tiny microbes. Fungi are an important part of these ecosystems because they help break down materials and can affect the health of animals like frogs. However, we still know very little about how fungi respond to urban environments. In this study, we looked at fungi living in water, sediment, and frog guts along a range of places from natural to highly developed areas in Worcester, Massachusetts. We found that some fungi were more common in cleaner, less developed sites, while others were more common in urban ones. These patterns suggest that fungi could be useful indicators of environmental health. Our work helps scientists better understand how urbanization affects nature and offers a new way to monitor and protect freshwater ecosystems.

## INTRODUCTION

Urbanization alters biodiversity and ecosystem function, affecting both macroscopic and microscopic life. Freshwater biodiversity is crucial for maintaining water quality and nutrient cycling, but urbanization degrades these systems by increasing pollution, habitat fragmentation, and impervious surface areas (ISA), which intensify runoff and pollutant loads (1–4). Additionally, urban discharge introduces novel microbial taxa, altering community composition and potentially reshaping ecosystem functions (5). ISAs, such as buildings and pavement, contribute to environmental degradation by intensifying heat island effects and increasing runoff elevating sediment loads (6). Stormwater and wastewater further introduce excess nutrients, heavy metals, and pharmaceuticals, degrading water quality (5, 7).

Urbanization negatively impacts waterway-associated organisms, including amphibians, which serve as environmental indicators due to their sensitivity to habitat changes. Habitat loss is a major driver of global amphibian declines (8, 9), while pollution and habitat modification also disrupt microbial communities, including those associated with amphibians (5, 10). As amphibians persist in varying degrees of degraded environments, shifts in their microbiomes serve as signals of ecosystem health before extreme habitat loss occurs. While the effects of urbanization on amphibians have been documented (11, 12), urban-driven changes also extend to microbial communities, which is comparatively less well studied. Despite the generally cryptic nature of the members of a microbial community, their composition is essential to overall ecosystem function.

Microorganisms play essential roles in nutrient and energy cycling and exhibit rapid growth, making them highly sensitive to environmental changes (10, 13, 14). As a result, microbial taxa can serve as potential biological indicators of environmental and host health. Microbiomes are the community of microbes comprised by viruses, bacteria, archaea and microscopic eukaryotes. The microbiomes of gastrointestinal tracts are particularly critical for host physiology, immune function, and behavior, influencing the response to environmental shifts (14).

Despite their ecological importance, fungi have been historically overlooked in microbiome studies, even though they play key roles in host metabolism, carbohydrate degradation, pathogen resistance, and immune regulation (14–16). Understanding fungal communities (the “mycobiome”) is essential, as host-fungal interactions can support amphibian health or contribute to population declines, especially in urban environments where microbial diversity is reshaped by environmental pressures (14). Microbiome research provides valuable insights into amphibian conservation by revealing how urbanization affects microbial communities, host health, and ecosystem stability [12–14]. While most urbanization studies on microbial communities have focused on bacteria (2, 6, 17, 18), recent research highlights fungi and the mycobiome as potential environmental health indicators(19–21). However, prior microbiome studies typically examine urban impacts on only a single habitat type, such as water, sediment, or host-associated communities (22–24).

Here, we studied the shifting biodiversity and ecology of the mycobiome along an urbanization gradient in Worcester, MA, across water, sediment, and frog-associated microbiomes. Worcester is the second most populous city in New England (USA) and has a long history as a manufacturing hub, with high levels of human activity and pollution from goods production factories. Across the city, there are numerous points of contacts with these historical production centers that directly connect with many waterways that converge into the Blackstone river (25). The northernmost parts of Worcester have lower population density and larger areas of protected land, as well as freshwater reservoirs. The Tatnuck Brook watershed presents a gradient of rural, more protected areas in the north to increasingly developed areas to the south, making it ideal to study the impact of urbanization on freshwater ecosystems.

We tested the hypothesis that increasing urbanization alters fungal community structure in the waterway, leading to shifts in diversity across habitats. Specifically, we predicted that urbanization would drive changes in community composition, with certain fungal taxa increasing in abundance in highly urbanized sites while others decline, reflecting environmental filtering along the urbanization gradient. We focused on accomplishing two main objectives: 1) To elucidate the differences of the fungal microbial communities in freshwater ecosystems across the microbiomes of sediment, water and frog gastrointestinal tracts, and 2) to characterize potential changes in taxonomic and functional fungal microbial diversity of these communities associated with a rural to urban gradient.

## METHODS

### Study Site Selection

We collected samples from three main habitats: fecal pellets from three frog species, sediment cores, and water samples across the Tatnuck Brook watershed system (Figure 1). Ten sites spanning approximately 10 km from north to south were sampled. Sites were a mix of ponds and streams that varied strongly in their exposure to human activity. Percent impervious surface area (PISA) was calculated for each site to determine urbanization levels using QGIS V 3.30.1 (26). The impervious surface area (ISA) layer, created by data collected in 2005, was extracted from publicly available MassGIS data (https://www.mass.gov/info-details/massgis-data-impervious-surface-2005). Urbanization levels were classified by the PISA value for each site, characterized as low (<10%), medium (10–30%), and high (>30%) urbanization categories as determined by Hincher et al.(27).

**Fig 1.**
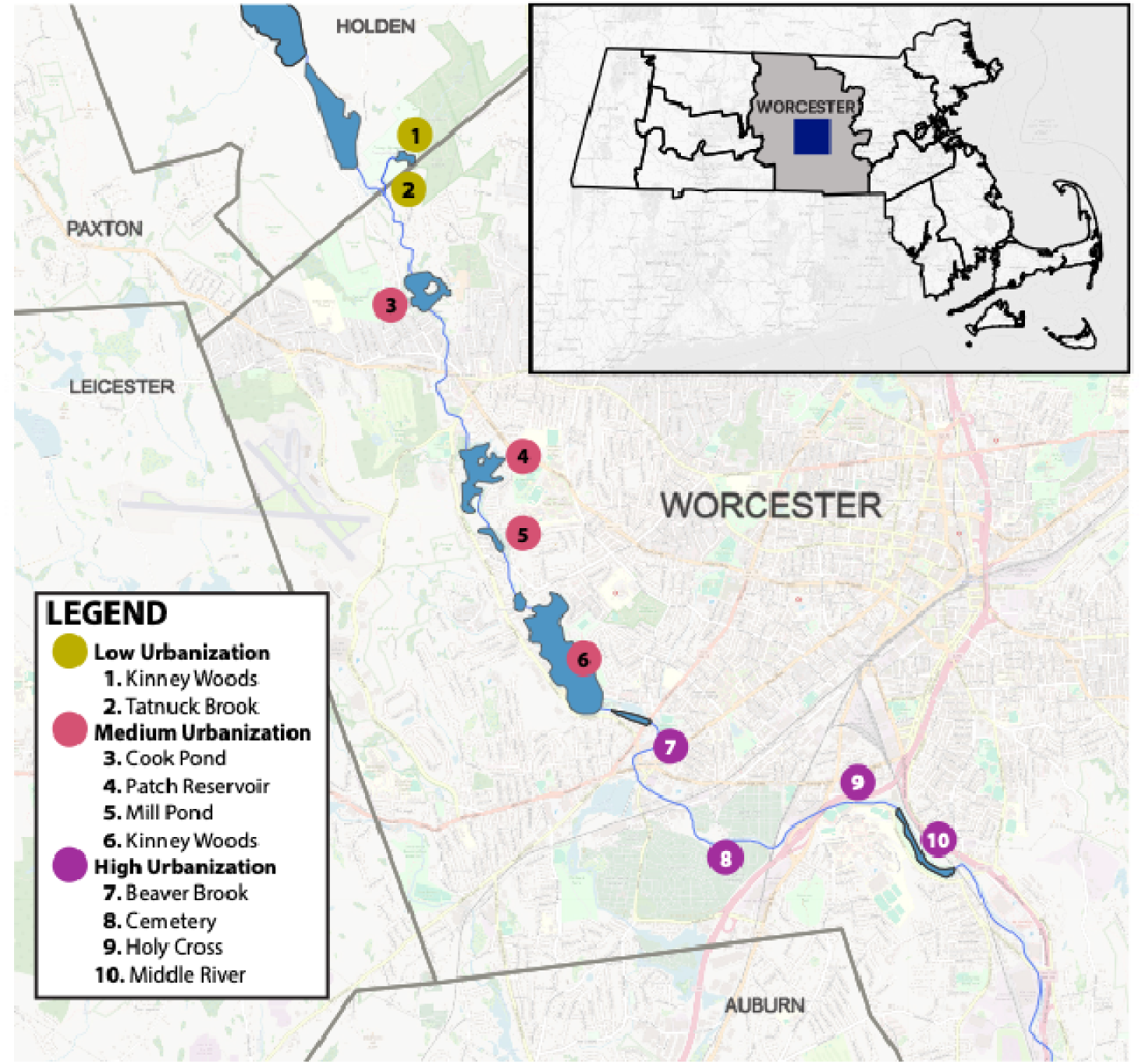
Map of the sampling sites along the Tatnuck Brook waterway in Worcester, Massachusetts. The y-axis represents longitude, and the x-axis represents latitude. Circles depict the location sampling sites, color-coded by urbanization level, and corresponding site names are listed in the legend.

Three frog species were collected: *Lithobates catesbeianus* (American bullfrog)*, Lithobates clamitans* (Green frog), and *Lithobates palustris* (Pickerel frog). These are common local frog species and of no conservation concern and thus were selected for non-destructive capture and fecal pellet isolation. All amphibians were captured at sites authorized by the Massachusetts Division of Fisheries and Wildlife (Permit 16.23SCRA). The capture-release protocol was approved by Clark University’s Institutional Animal Care and Use Committee (IACUC, Protocol 037R). Frogs were temporarily housed in sterile collection bags until defecation occurred, after which they were released at their original capture site. Fecal samples were then collected using a sterile scoopula. The fecal sample was covered with 20% glycerol and stored at −20 °C until DNA extraction.

Sediment core samples were taken at stream sites using a Large Universal Corer (Aquatic Research Instruments, Idaho, USA) to a depth of ∼5 cm at four locations randomly chosen at each site and kept separate. After sampling, each core was homogenized and ∼50 ml duplicate subsamples were stored separately of each core at −80 °C until DNA extraction.

Water samples were collected using acid-washed 4 L cubitainers, rinsed using water downstream of the collection point. Water samples were placed on ice until filtered later that day. Water samples were filtered by peristaltic pump by drawing water through a 25 µm Nitex mesh covering the intake opening of the peristaltic tubing and then passing through a 47 mm 3 µm polycarbonate filter (Cat # PCT3047100, Sterlitech, USA) within a stainless-steel filter holder to collect eukaryotic cells. The 3 µm filters were then aseptically removed and stored in tubes kept at −80 °C until DNA extraction.

### Environmental DNA Extraction

Fecal DNA was extracted from frog fecal samples and sediment samples using the QiAmp PowerSoil Pro DNA Kit (Cat # 47021, Qiagen, Maryland, USA). For water DNA extraction, a phenol chloroform protocol was used to extract DNA from filters. Modifications to the protocols can be found in the Supplementary Materials. DNA concentrations of all samples were determined using a Qubit 4 Fluorometer with a dsDNA High Sensitivity kit (ThermoFisher Scientific, Massachusetts, USA). The purity was measured with a Nanodrop One Microvolume UV-Vis Spectrophotometer (ThermoFisher Scientific, Massachusetts, USA).

### ITS1 Metabarcode sequencing and analysis

The ITS1 region was amplified using the standardized primer set in Bellemain et al. (2010) to conduct metabarcode community analysis of fungal communities. Amplification, library preparation, and 2X250 PE sequencing was done by NovoGene Inc. (Sacramento, CA) using on an Illumuna NovaSeq6000 sequencer. Illumina adapter sequences were removed, and sequences were demultiplexed by NovoGene. The resulting raw reads were quality trimmed with cutadapt (v3.4) (28). Downstream processing of trimmed reads was performed using DADA2 (v1.16) (29), and taxonomy was assigned to resulting amplicon sequence variants (ASVs) using the general Fungi UNITE database (version 9.0) with a naïve Bayesian classifier and a minimum bootstrap confidence threshold of 80% (30).

### Fungal Community Analysis

All data analyses were conducted in R (4.4.2) (31) and RStudio (32). Phyloseq (V 1.44.0) (33) was used for microbiome data processing such as deduplicating reads, defining ASVs and generating counts per ASV. Relative abundances were calculated for downstream analysis. Rarefaction curves were generated with the function rarecurve from vegan (v. 2.6) (34). Reads were not rarified for the subsequent analyzes as this type of data stratification tends to remove rare, but important taxa (35).

For alpha-diversity analyzes, Shannon and Simpson diversity indices were computed with the diversity function in vegan (34). Analysis of Variance (ANOVA) was performed to analyze differences in alpha diversity, using Shannon or Simpson’s indices as response variable, and habitat and urbanization levels as the explanatory variable. Assumptions for ANOVA were checked using Q-Q plots for normality and Bartlett’s test for homogeneity of variance. Post hoc analyses for significant ANOVA results were conducted using Tukey’s HSD tests. Boxplots were generated using ggplot2 (36).

Beta diversity was compared using permutational multivariate ANOVA (PERMANOVA) using the adonis2 function in the vegan package (37) with 1,000 permutations. Nonmetric multidimensional scaling ordination (NMDS) plots were generated using Bray-Curtis dissimilarity values. For the PERMANOVA, the Bray-Curtis dissimilarity values were used as the response variables, and the habitat or urbanization levels as the categorical explanatory variables. For the frog fecal samples, we also included the frog host species and the interaction of urbanization level and host species as the categorical variables. Post hoc pairwise PERMANOVA analyses were conducted to determine which specific variables exhibited significant community differences from each other using the pairwiseAdonis package (v. 0.4.1) (38) with Benjamini-Hochberg (BH) p-value correction for multiple test comparisons (39). The top 20 most abundant taxa per habitat were ranked by mean relative abundance.

Differential abundance analyses were conducted using DESeq2 (40) to identify taxa with significant changes in absolute abundance per source across urbanization levels. Variance-stabilizing transformation (VST) was applied to normalize count data. Differentially abundant taxa between urbanization categories were identified using the DESeq function, with a significance threshold of adjusted p-value < 0.05. Log2 fold changes were computed, where an ASV with log2FC ≥ 5 or ≤ −5 indicates a significant presence in one condition compared to the other.

### Fungal trophic guild abundances

Fungal ecological guild information was extracted using FUNGuild (v. 1.1) (41) and its associated database, with ASVs without annotation being removed from further analysis. After assignment of guild to each ASV, analysis targeted ASVs assigned to only one trophic guild with a confidence ranking of ‘probable’ or ‘highly probable.’ Significant differences in the absolute abundance of each trophic mode were analyzed using a Kruskal-Wallis test in R, after assumptions for ANOVA were checked and violated (Q-Q plots for normality and Bartlett’s test for homogeneity of variance) as well as a permutation ANOVA using the aovp function of the lmPerm R package using 5,000 permutations (42) and pairwise permutation tests in the pairwisePermutationTest function from the rcompanion R package using 10,000 permutations and the false discovery rate (FDR) correction (43). For each guild (Saprotroph, pathogen and symbiont), the abundance was used as the response variable and the urbanization category as the explanatory variable) for both Kruskal-Wallis test and permutation ANOVA.

## RESULTS

### Sampling and Meta-amplicon Sequencing

Water (planktonic), stream sediment, and frog fecal samples were collected from the Tatnuck Brook waterway, spanning a north-to-south urbanization gradient in Worcester (Fig. 1 (Table 1). The northernmost sites—Kinney Woods (5.1%) and Tatnuck Brook (9.0%)—were classified as low urbanization. The next four sites had medium urbanization (17.2–25.6%), while the five most southern, downstream sites exceeded 45% PISA, with Holy Cross (56.7%) and Beaver Brook (56.6%) showing the highest urbanization levels (Table 1).. This 10 km gradient reflects a transition from rural to developed regions.

**Table 1.** Classification of sampling sites in the Tatnuck Brook watershed based on percent impervious surface area (PISA)

| Site Name | Site ID | Percent Standard Impervious Surface Area (PISA) | Urbanization Category |
| --- | --- | --- | --- |
| Kinney Woods | KP | 5.1% | Low |
| Tatnuck Brook | PRS, CP2 | 9.0% | Low |
| Cook Pond | Cook | 17.2% | Medium |
| Patch Reservoir | PR | 22.0% | Medium |
| Mill Pond | MP | 25.6% | Medium |
| Coes Pond | Coes, CR2, CR3 | 24.7% | Medium |
| Beaver Brook | BB1 | 56.6% | High |
|  | BB2 | 48.1% | High |
|  | BB3 | 52% | High |
| Cemetery | MR1 | 46.2% | High |
| Holy Cross | MB | 56.7% | High |
| Middle River | MR, MR2 | 51.8% | High |

A total of 129 microbiome samples were collected, with 112 yielding high-quality sequencing data (fecal n=60, sediment n=30, water n=22) (Supp. Table 1). These samples generated 25,608,648 paired-end raw reads, resulting in 121,368 unique amplicon sequence variants (ASVs) spanning 20 phyla, 80 classes, 214 orders, 596 families, 1,740 genera, and 2,879 species (Table 2). Rarefaction curves plateaued, confirming sufficient sequencing depth to capture taxonomic diversity (Supplementary Figure 1).

**Table 2.** Number of fungal ASVs assigned to each phylum across fecal, water, and sediment habitats. The table presents the number of ASVs associated with each fungal phylum (identified using the UNITE database) across the three sample habitats. Notably, *Ascomycota* and *Basidiomycota* dominate all habitats, while less common phyla such as *Rozellomycota*, *Chytridiomycota*, and *Mortierellomycota* show variable representation. The presence of rare phyla, including *Neocallimastigomycota* and *Entomophthoromycota*, highlights the diversity and complexity of fungal communities in these ecosystems

| Phyla<br>(UNITE) | Fecal | Water | Sediment |
| --- | --- | --- | --- |
| Aphelidiomycota | 15 | 9 | 15 |
| Ascomycota | 6313 | 1532 | 6580 |
| Basidiobolomycota | 130 | 35 | 48 |
| Basidiomycota | 2087 | 364 | 3812 |
| Blastocladiomycota | 14 | 9 | 53 |
| Chytridiomycota | 944 | 943 | 3506 |
| Entomophthoromycota | 1 | 0 | 5 |
| Glomeromycota | 11 | 7 | 42 |
| GS01 | 18 | 6 | 16 |
| Kickxellomycota | 12 | 0 | 11 |
| Monoblepharomycota | 8 | 16 | 24 |
| Mortierellomycota | 189 | 24 | 192 |
| Mucoromycota | 342 | 8 | 129 |
| Neocallimastigomycota | 0 | 0 | 5 |
| Olpidiomycota | 29 | 13 | 30 |
| Rozellomycota | 602 | 189 | 771 |
| Zoopagomycota | 9 | 6 | 8 |

### Alpha and Beta diversity of Fungal Communities Across Freshwater Habitats

Alpha and beta diversity analyses revealed significant differences in fungal communities across habitats and urbanization levels. Using Shannon’s diversity index, sediment samples were significantly more diverse than fecal and water samples (ANOVA, F = 37.95, p<0.001; Tukey’s post hoc, p<0.001), while fecal and water samples did not differ (Tukey’s post hoc, p = 0.132) (Figure 2). In contrast, there were no significant differences across habitats for Simpson’s diversity index (ANOVA, F = 1.39, p=0.253), suggesting similar dominance patterns among sources (Supplementary Figure 2a). Beta diversity analysis (PERMANOVA) confirmed distinct microbial communities across habitats (F = 5.79, p = 0.001, R² = 0.095), with pairwise comparisons showing significant differences between all groups (p = 0.001). Urbanization significantly influenced fungal community composition (F = 3.32, p = 0.001, R² = 0.050), though habitat type explained more variance, indicating a stronger role in shaping community structure. Fecal samples were dominated by *Basidiobolus* and *Cladosporium*, while sediments were dominated by *Lemonniera* (Supplementary Figure 3). *Only Candida* was consistently found in all habitats (Supplementary Table 2 and Supplementary Figure 3).

**Fig. 2.**
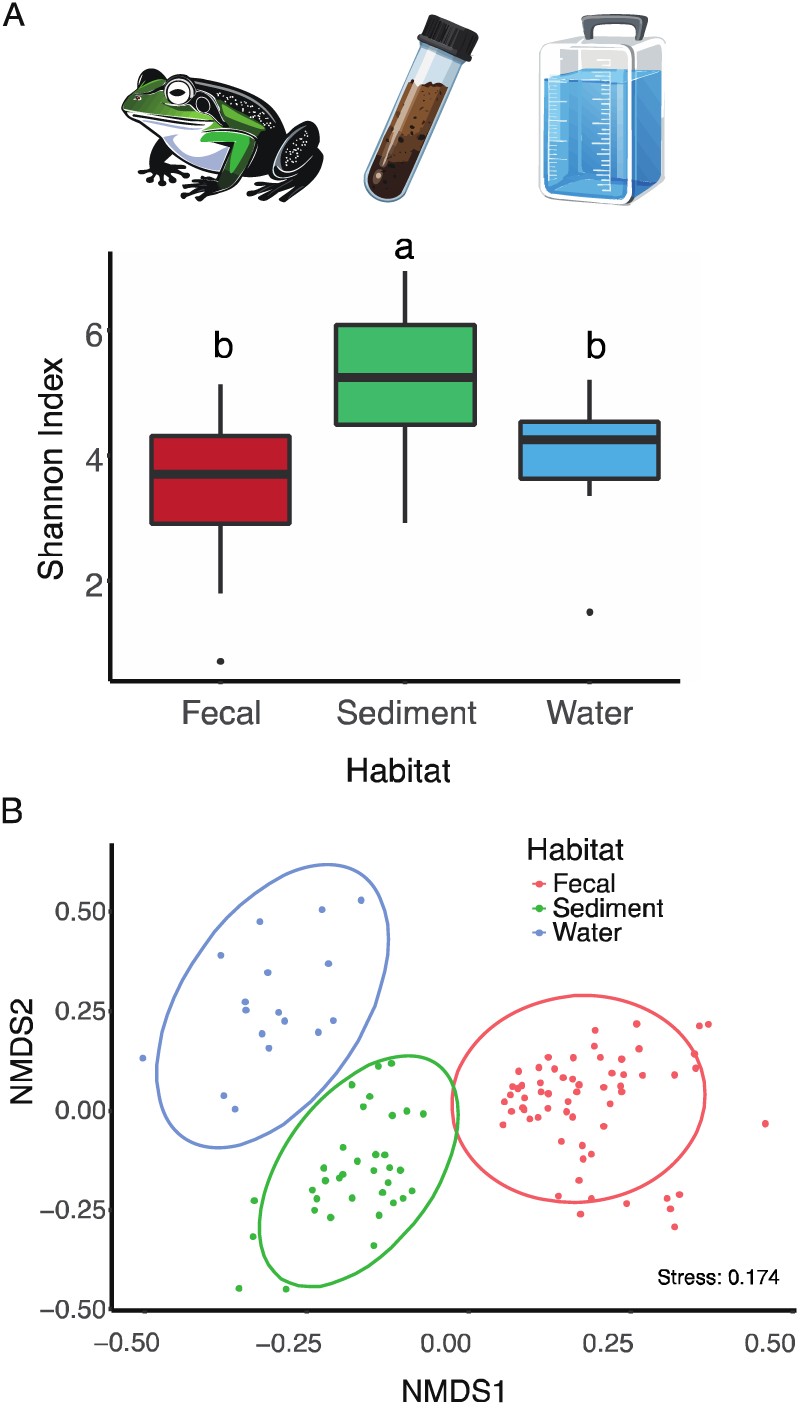
Diversity and community composition of microfungi associated with amphibian fecal pellets, sediment, and water samples. (a) Shannon diversity index of microfungal communities across three habitats: fecal pellets (Represented by a Green Frog), sediment (Represented by a soil sample in a tube), and water (Represented by a cubitainer). Letters above bars indicate statistical similarity or differences, with shared letters representing no significant difference (e.g., “a”). (b) Non-metric multidimensional scaling (NMDS) plot illustrating differences in microfungal community composition among the three habitats. Each point represents a sample, and its position reflects similarity in community composition. Axes represent the first two dimensions of ordination space (NMDS1 and NMDS2). Ellipses represent 95% confidence intervals for the multivariate t-distribution of the ordinated samples. All illustrations were generated in Adobe Illustrator.

### Effects of Urbanization on Fungal Communities

#### Fecal Fungal Communities

To control for the effect of habitat, samples were analyzed separately by habitat type to further test for relationships between community diversity and urbanization. In frog fecal samples, urbanization did not significantly impact alpha diversity using either Shannon’s diversity (*F* = 1.804, *p* = 0.174) or Simpson’s diversity (*F* = 0.01, *p* = 0.994). There was also no effect of frog species (Shannon: *F* = 0.502, *p* = 0.608, Simpson: *F* = 0.546, *p* = 0.608) or the interaction of frog species and PISA categories (Shannon: *F* = 0.507, *p* = 0.605, Simpson: *F* = 0.585, *p* = 0.560).

However, although there were no differences in alpha diversity, there were significant differences in fungal community composition between frog species (F=1.29, p=0.020, R²=0.038) (Supplementary Figure 2b) and urbanization levels (F=3.88, p=0.001, R²=0.116) (Figure 3a), with the latter having a stronger effect. The interaction between host species and urbanization was not significant (F=1.091, p=0.169, R²=0.032). Pairwise PERMANOVA tests confirmed significant community composition differences across urbanization levels (p=0.001 for all comparisons). Notably, *Basidiobolus* and *Candida* were abundant in low urbanization, while *Botrytis*, *Alternaria*, and *Parasarocladium* were more common in highly urbanized sites (Supplementary Table 2). Differential abundance analysis identified 125 taxa showing significant differences between low and high urbanization, with 82 favoring low and 43 favoring high urbanization (Table 3, Supplementary File 1).

**Fig. 3.**
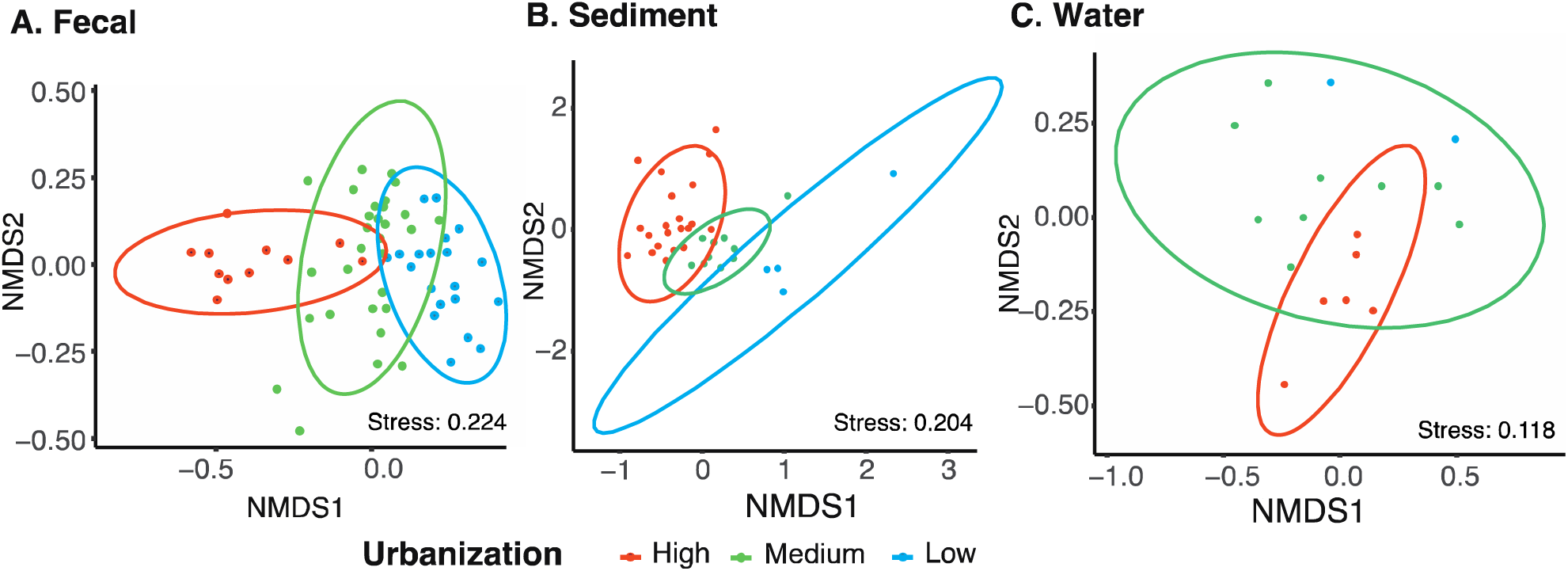
Effects of urbanization on microfungal community composition in amphibian fecal pellets, sediment, and water samples. NMDS plots show the ordination of microfungal communities with respect to different levels of urbanization (High, Medium, Low) for samples segregated by habitat type: a) frog fecal samples b) stream sediment samples, and c) planktonic water samples. For each figure the axes represent the first two dimensions of ordination space (NMDS1 and NMDS2). Each point represents a sample, which are color coded by urbanization level. Ellipses represent 95% confidence intervals for the multivariate t-distribution of the ordinated samples. Only two low-urbanization water samples were generated (c) so an ellipse could not be drawn for that sample subset.

**Table 3.**
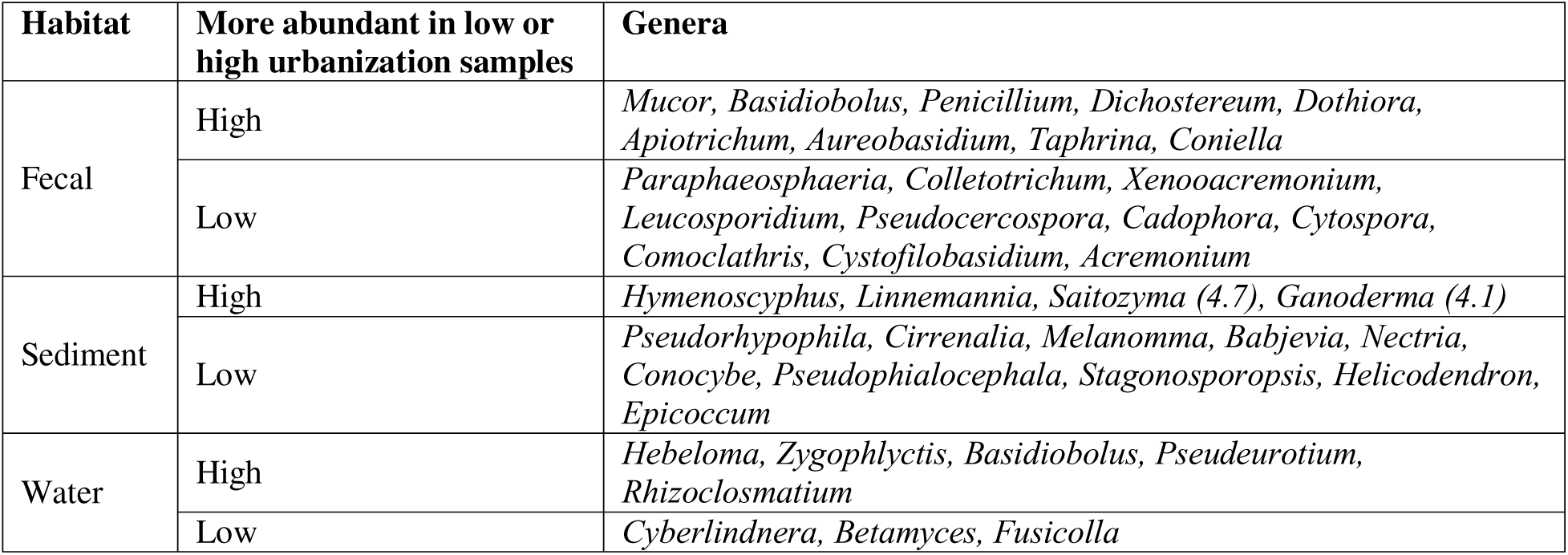
Fungal genera significantly enriched in low or high urbanization samples across different habitats, based on DESeq2 analysis. In most cases, genera were detected in one level but not the other, but otherwise the log2-fold difference is listed in parentheses after the genus name.

#### Sediment Fungal Communities

Sediment samples exhibited no significant difference in Shannon’s (*F* = 2.16, *p* = 0.132) or Simpson’s (*F* = 1.60, *p* = 0.217) diversity indices across urbanization levels. However, beta diversity was significantly influenced by urbanization (PERMANOVA, *F* = 2.32, *p* = 0.001), in this case explaining 13% of the observed variation (Figure 3b). Post hoc pairwise PERMANOVA confirmed significant differences across all urbanization levels as well (p=0.001 for all comparisons). Low urbanization sites had unique genera like *Apodus* and *Podospora*, while high urbanization sites were characterized by *Neoascochyta* and *Candida* (Supplementary Table 2). Out of 33 taxa that were differentially abundant between low and high urbanization, ten and twenty-three were more common at low and high urbanization, respectively (Table 3, Supplementary File 2)

#### Planktonic Fungal Communities

As with the other habitats, planktonic fungal communities showed no significant change in Shannon’s (*F* = 0.24, *p* = 0.787) or Simpson’s diversity indices (*F* = 0.32, *p* = 0.729) with urbanization. Nevertheless, community composition was significantly structured by urbanization (PERMANOVA, *F* = 1.33, *p* = 0.003, R^2^ = 0.159) (Figure 3c). 16% of sample variance was explained by urbanization, again notably lower than for frog fecal mycobiomes. There was overlap in the clustering of medium and high urbanization samples, but PERMANOVA shows a significant difference in community composition between these groups (F = 1.528, p = 0.006, R^2^ = 0.105). There were only two low-urbanization water samples, so we were unable to test for significant differences in community composition of these sites from other urbanization sites.

The top 20 genera analysis showed that low and medium urbanization sites had greater abundances of *Basidiobolus*, *Clavispora*, and *Zygophlyctis*, while high urbanization sites featured *Aureobasidium* and *Metarhizium* (Supplementary Table 2). We identified 95 differentially abundant taxa between low and high urbanization sites. Chytridiomycota ASV’s represented the most differentially abundant taxa, with 45 ASV’s represented in low urbanization, in comparison to 13 at higher urbanization (Table 3, Supplementary File 3).

#### Fungal Trophic Mode Analysis

A total of 35,854 fungal ASVs were classified into pathotrophs, saprotrophs, symbiotrophs, and unknowns, with 58% of the ASVs labeled as unknown due to multiple assignments or low confidence rankings (Figure 4). While most Kruskal-Wallis and permutation-based ANOVA tests did not detect significant differences in trophic mode abundance across urbanization levels (Supplementary Tables 3 and 4), the saprotroph community in water samples showed a significant effect of urbanization in the global permutation ANOVA (p = 0.036). However, post hoc pairwise permutation tests did not reveal

**Fig. 4.**
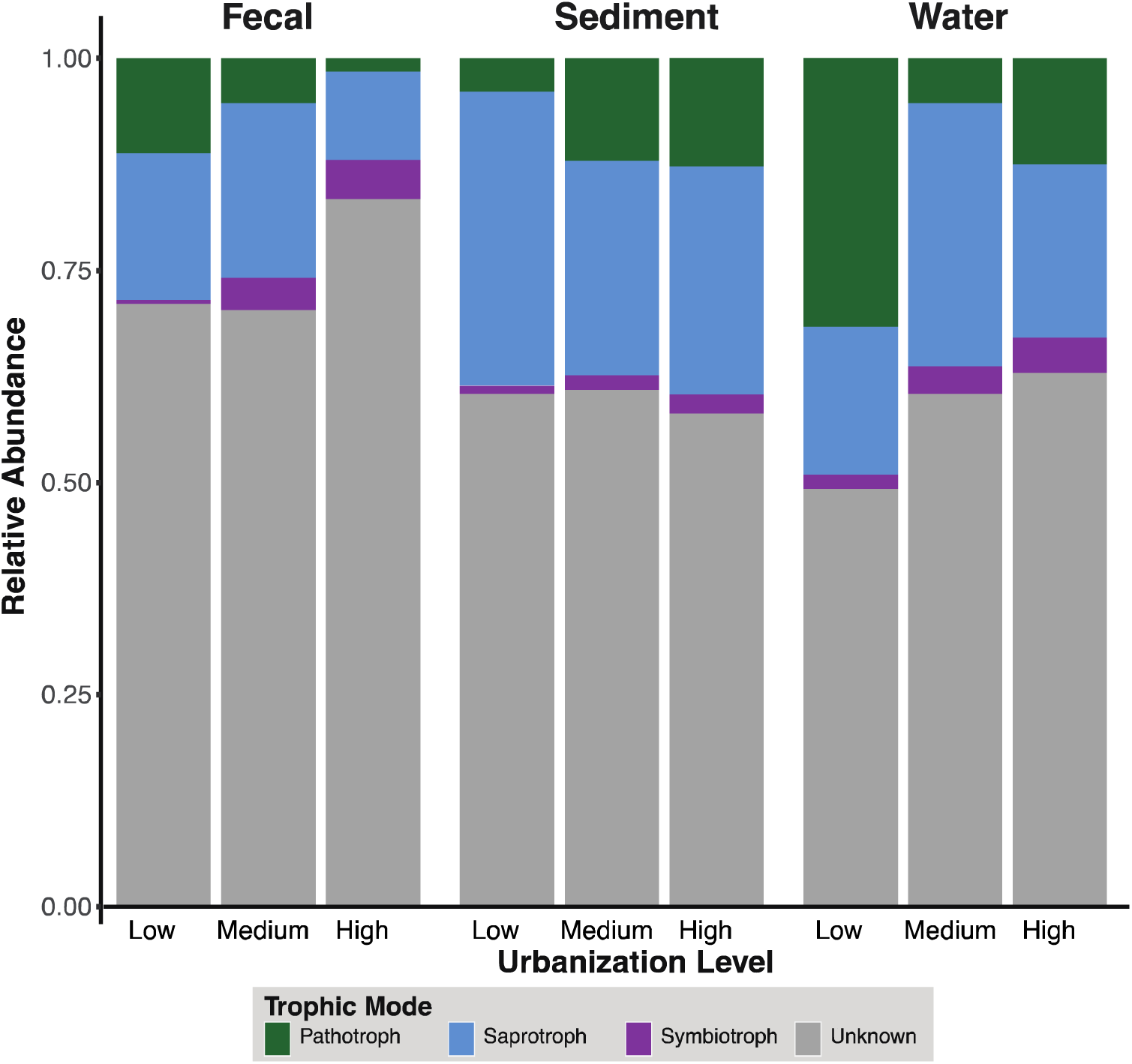
Proportional representation of fungal trophic modes across fecal, sediment, and water samples under varying urbanization levels. The bar plots illustrate the relative abundance of fungal trophic modes (pathotrophs, saprotrophs, and symbiotrophs) and unknowns according to the number of ASVs assigned to each in samples collected from fecal pellets, sediment, and water. Urbanization levels are compared within each sample type, highlighting shifts in trophic mode proportions in response to urbanization gradients.

significant differences between specific urbanization categories after false discovery rate (FDR) correction, despite a marginal trend between high and medium sites and low and medium size (adjusted p = 0.16 for both paiwise comparisons). These findings suggest that while saprotrophic communities in water are influenced by urbanization at a broad level, the directional differences between individual urbanization categories are subtle and may reflect gradual community shifts rather than discrete stepwise transitions. Such shifts may still carry ecological implications for nutrient cycling and organic matter processing in urban-impacted aquatic systems.

## DISCUSSION

Urbanization is reshaping ecosystems, altering both macroscopic biodiversity and microbial communities. While urban impacts on bacterial and archaeal communities have been studied, fungal communities remain underexplored. Primary, previous research has often focused on a single habitat, be it environmental (44–46) or host associated (47–50) and less frequently in aquatic environments. This study examines three interconnected microbial environments, including both environmental and host derived sources. Our results show that while alpha diversity remained stable, beta diversity shifted significantly across urbanization level, indicating taxonomic turnover rather than biodiversity loss. This aligns with previous studies showing that urbanization reshapes microbial composition without reducing species richness (51–53).

There were some notable differences in strength of correlation of urbanization with community composition. Urbanization influenced fungal community composition across habitats, but habitat type was a stronger predictor (9.5% vs. 6%) when considering all samples together. This is not surprising because habitat type is a strong driver of prokaryotic community similarity (54). When considering each habitat separately, urbanization explained similar proportions of variance in community composition (12%, 13%, and 16% for fecal, sediment, and water habitats respectively). Within fecal samples, host species identity explained only 4% of variance, suggesting that urbanization exerts a stronger influence on microbial turnover than interspecies differences. These findings indicate urbanization has broad-sweeping impacts of altering fungal communities across various components of the aquatic ecosystem.

The observed shifts in fungal community composition suggest that urban environments act as ecological filters, favoring taxa adapted to pollutants, altered nutrient dynamics, and habitat disturbances (22–24, 47, 55, 56). However, the specific mechanisms driving these shifts remain unclear. Future metagenomic studies could identify key genes and pathways involved. Additionally, the implications for amphibian health and water quality require further investigation. By analyzing fungal taxa across freshwater habitats, our findings highlight how urbanization shapes microbial diversity and underscore the need for research into the functional roles of urban-adapted fungi in host-microbe interactions, biogeochemical cycling, and ecosystem stability.

### Differences in Fungal Community Responses Across Sources

The three primary habitats—sediment, water, and host associated fecal matter—demonstrate distinct community compositions influenced by urbanization, reflecting the diverse ecological niches within the watershed. Sediment, with its high fungal diversity, supports saprotrophic fungi crucial for organic matter breakdown, likely due to nutrient-rich microenvironments that favor fungal growth and metabolic diversity [40, 41, 42]. The species richness in sediment may also stem from its role as a microbial sink, capturing inputs from water and detritus, and supporting taxa that interact with organic substrates.

The planktonic communities exhibited fewer taxa but showed notable resilience to urbanization-induced changes, particularly in community composition. The persistence of certain genera, like *Clavispora* and *Aureobasidium*, across urbanization levels in water suggests their functional adaptation to the aquatic environment, highlighting their ecological relevance as nutrient recyclers and decomposers in freshwater systems (60). The resilience observed may be attributed to the natural flow dynamics in water bodies, which might buffer against certain urban pollutants (61, 62).

Frog fecal samples showed the greatest compositional shifts with urbanization. *Basidiobolus*, a genus linked to amphibian gut health, was abundant in low urbanization sites but absent in high urbanization samples, suggesting sensitivity to urban stressors. Previous studies found *Basidiobolus* to be a dominant gut fungus, representing 84% of gut fungi across various salamander species, 29% of gut fungi in various species of frogs (63), and comprising 60% of gut fungi in slimy salamanders (*Plethodon* spp.) (64). Despite its prevalence in amphibians, in this study it was abundant only in low urbanization sites. This suggests *Basidiobolus* may be a useful bioindicator for habitat quality and amphibian-microbiome interactions.

### Novel discoveries in fungal ecology

The *Rozellomycota* were highly represented across all sample types and mostly in fecal and sediment samples. They are typically cryptic or parasitic, so they are often linked to other fungi or protists in soil. The presence of *Rozellomycota* in both amphibian fecal pellets and sediment expands their known habitat range. Their detection in aquatic environments suggests potential roles in organic matter decomposition, symbiosis, or host-microbe interactions that warrant further study.

Additionally, this study revealed the presence of *Basidiobolus* within the planktonic water communities, marking one of the first records of this genus in freshwater systems. This genus is typically associated with amphibian gastrointestinal tracts or terrestrial environments such as leaf litter, so its presence in water possibly suggests environmental transmission pathways that extend beyond known host associations and a broader ecological versatility (65–67). These findings suggest fungal adaptability in urban-impacted freshwater systems, with urbanization potentially facilitating the movement of terrestrial taxa into aquatic and host-associated environments. The presence of *Rozellomycota* in novel habitats highlights the need for further studies using multi-marker approaches to clarify their ecological roles, diversity, and contributions to nutrient cycling and ecosystem function.

### Implications of Trophic Mode Shifts in Urbanized Environments

Trophic mode analysis provides insights into how urbanization may influence the ecological roles of fungal communities. While no significant differences in mode were found with urbanization level, this apparent stability may reflect ecological resilience rather than a lack of response. The consistency in saprotrophs, pathotrophs, and symbiotrophs suggests that key ecosystem functions are maintained despite shifts in community composition. This points to possible functional redundancy within fungal communities, allowing them to sustain roles like decomposition and symbiosis even under urban stress. Other trends, such as saprotrophic fungi in water samples exhibited a significant effect of urbanization in the global permutation ANOVA (p = 0.036), although follow-up pairwise permutation tests did not yield statistically significant differences between specific urbanization categories after FDR correction. This pattern suggests a broad but gradual shift in saprotrophic composition in response to increasing urban impact, potentially linked to altered nutrient regimes or detrital inputs in urban waterways. In parallel, we observed non-significant but consistent trends of increasing pathotroph abundance in sediment and planktonic habitats with higher urbanization, echoing patterns reported in other studies (68, 69). This shift could impact amphibians and aquatic ecosystems by increasing pathogen loads, contributing to biodiversity loss. Increased urban organic waste may support more opportunistic fungi, while reduced plant diversity often seen with urbanization likely limits symbiotic fungi. The heightened pathotroph abundance in urbanized sediments and water raises concerns about potential amphibian health risks, as pathogenic fungi, such as chytrids, are known to have devastating effects on amphibian populations (70). However, the high proportion of unclassified ASVs underscores the need for further metagenomic research to clarify functional shifts in fungal communities along urban gradients.

### Urbanization as a Filter for Resilient or Opportunistic Fungal Taxa

The observed shifts in fungal communities align with the concept of biotic homogenization, where urbanization filters out sensitive taxa and selects for those resilient to anthropogenic pressures. This effect, where resilient taxa dominate urban ecosystems, highlights a trend toward microbial communities dominated by opportunistic or anthropophilic species (71–73). Such homogenization can reduce functional diversity within ecosystems, potentially diminishing the resilience of microbial communities to future environmental changes (71–73).

### Conservation and Restoration Implications

The distinct fungal community compositions across urbanization levels and sources in this study highlight the need for conservation and restoration efforts tailored to urbanized freshwater systems. The high fungal diversity observed in sediment, especially in low urbanization sites, suggests that sediment habitats act as refuges for microbial biodiversity. Preserving and restoring sediment-rich areas along urban watersheds could help maintain fungal diversity and associated ecosystem functions, such as nutrient cycling and organic matter decomposition.

These results also imply that some fungal taxa, particularly those in frog GI tracts, may be more sensitive to urban stressors and can be bioindicators for assessing environmental health in freshwater systems (74, 75). For instance, the absence of *Basidiobolus* in high urbanization fecal samples suggests it could serve as a sensitive indicator of habitat quality. Conservation strategies could include monitoring this genus across watersheds to identify areas of significant environmental degradation and inform mitigation actions.

## Conclusions

This study serves as an initial step toward understanding the effects of urbanization on fungal communities in freshwater ecosystems, and the interconnected nature they form across different trophic levels. Here we find consistent patterns of impact across three inter-connected microbial habitats in a freshwater ecosystem: stable alpha diversity but significant shifts in beta diversity associated with urbanization. Future research should expand upon these findings by integrating shotgun metagenomics to determine specific metabolic pathways affected by urbanization. Additionally, studies tracking temporal fungal communities could inform the responses of these communities to increasing urban development. This interconnectivity of freshwater systems renders it critical to study environmental microbial communities in tandem with host microbial communities since they serve as source pools for host microbiomes. Therefore, sediment and planktonic microbial communities in freshwater systems are also of interest as they not only perform vital ecosystem services but also contribute to the assembly of microbial communities in amphibians.

## Acknowledgements

Special thanks to Dr. Anne Devan Song for her field expertise and help with our collection of amphibians. This project was funded by the Clark University’s Academic Innovation Fund.

## Data availability

All raw sequencing data has been deposited in the Short Read Archive (SRA) under Bioproject number PRJNA1244802. All supplementary data, including all supplementary files, tables, figures, R objects and code can be downloaded from GitHub at (https://github.com/TabimaLab/Fungal_Watershed_Mycobiome).

