## Supplementary Materials for "Effects of urbanization on the microbial fungal communities from water, sediment, and amphibian hosts across a rural-to-urban waterway in Worcester, Massachusetts"

**SUPPLEMENTARY METHODS**

**DNA extraction from samples across multiple habitats:**

For fecal DNA extraction, the manufacturer’s protocol was modified by: (1) if necessary, fecal samples were spun down and excess glycerol was discarded, (2) a solid piece of the fecal sample (approximately 250 mg) was placed directly into the manufacturer’s bead tube, (3) initial vortex time increased to 15 minutes, and (4) the C3 and C6 buffers were incubated at 65°C prior to use. Sediment DNA was extracted from sediment samples using the QiAmp PowerSoil Pro DNA Kit (Qiagen, Maryland, USA). The manufacturer’s protocol was modified by: (1) sediment samples were removed from the -80°C and thawed for 30 minutes, (2) sediment sample were vortexed to homogenize, (3) initial vortex time was increased to 15 minutes, and (4) the C3 and C6 buffers were incubated at 65 °C prior to use.

For water DNA extraction, the protocol used was modified from [24, 25]. The following modifications were made: (1) the DNA Extraction Buffer (DEB) was heated to 65°C in a water bath prior to use, (2) the DEB was then added directly into the tube with the filter (2) 10 µL Proteinase K was added along with the DEB, (3) all bead beating steps were skipped, (4) during the Ethanol precipitation step, one volume of 100% molecular-grade isopropanol was added instead of two volumes of 100% ethanol, (5) incubation with isopropanol at -20°C for 2 hours instead of overnight, and finally (6) the DNA pellet was resuspended in the elution buffer at 65°C for 10 minutes.

**SUPPLEMENTARY MATERIALS**

**Supplementary Fig. 1**

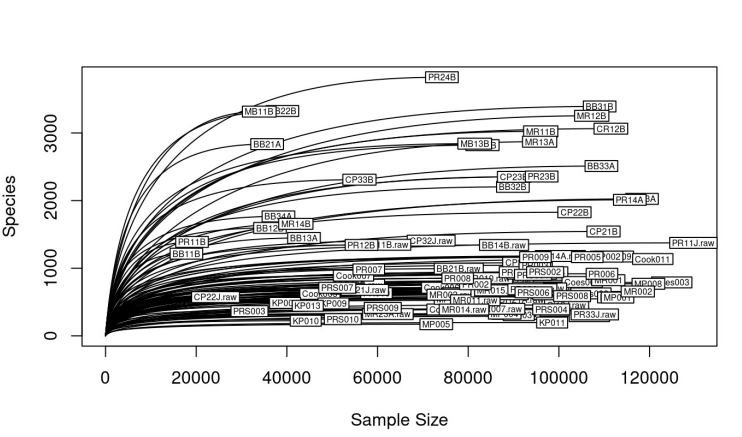

Rarefaction curves showing the observed fungal ASV richness across all samples. The curves illustrate the relationship between sequencing depth and the number of unique ASVs detected, indicating the adequacy of sampling effort. The curves begin to approach saturation, suggesting that sequencing depth was sufficient to capture most of the fungal diversity present.

**Supplementary Fig 2.** A). Boxplot representing Simpson diversity index of microfungal communities across three habitats: fecal pellets, sediment, and water. B). NDMS plot representing the beta diversity (using the Bray-Curtis dissimilarity metric) of microfungal communities on fecal pellets across three common frog species.

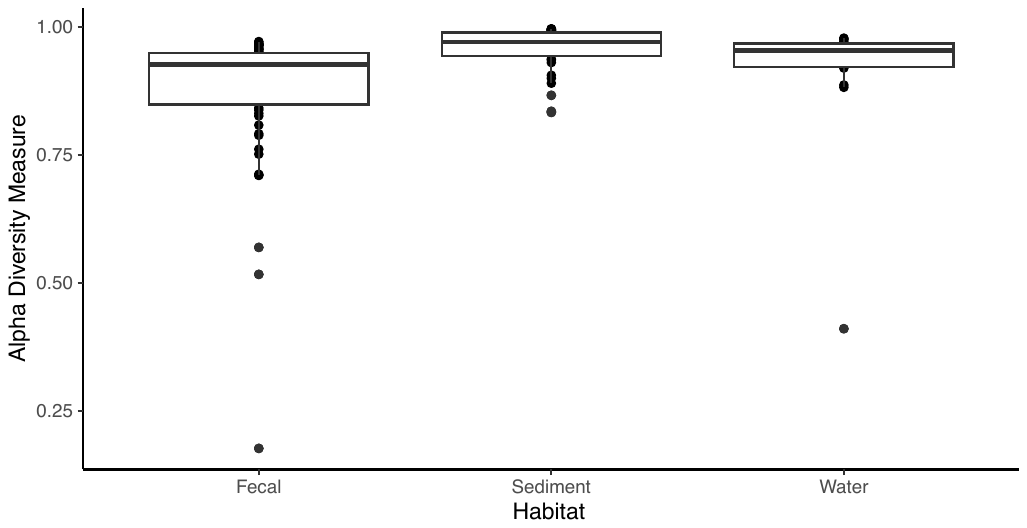

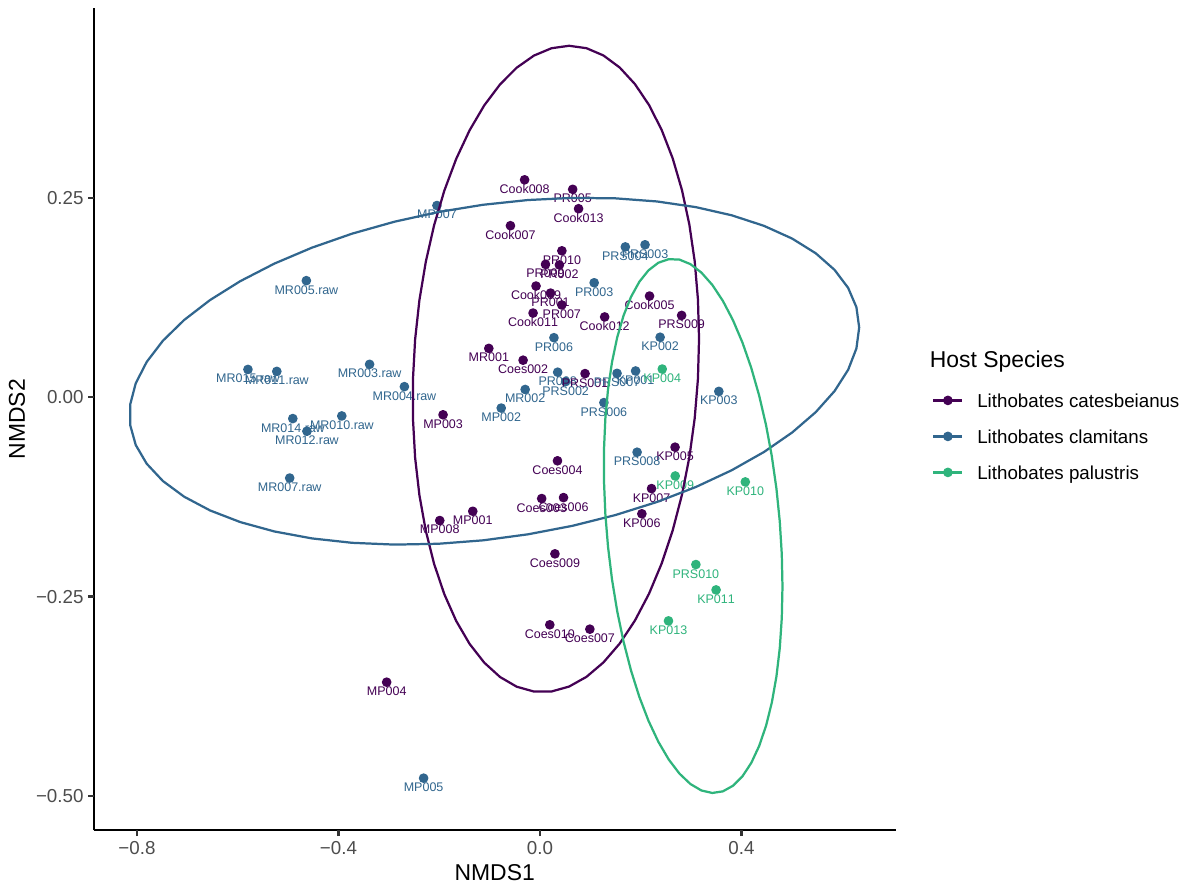

**Supplementary Fig 3.** Barplot representing the top 20 fungal genera across fecal, water, and sediment samples

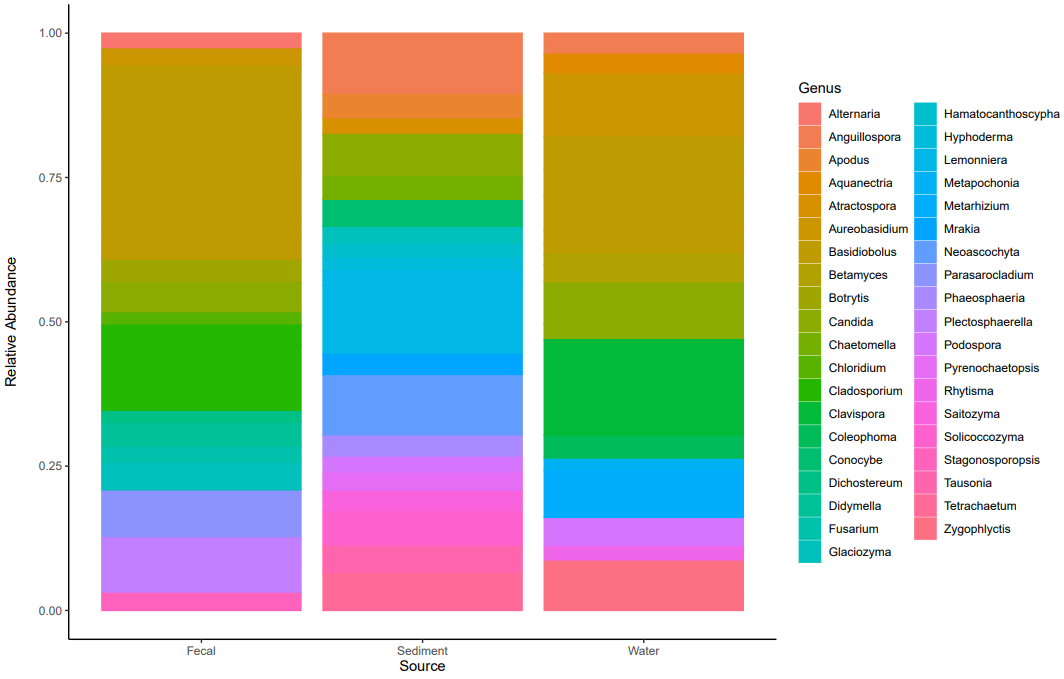

**Supplementary Tables**

**Supplementary Table 1**. Sample metadata for all collected fecal, water, and sediment samples, including site, habitat type, host species, and PISA classification.

| **Sample ID** | **Sites** | **Sample Type** | **Site Type** | **Urbanization Category** | **Host Species** | **PISA** |
| --- | --- | --- | --- | --- | --- | --- |
| BB11A.raw | Beaver Brook | Water | stream | High | <NA> | 56.6 |
| BB11B | Beaver Brook | Sediment | stream | High | <NA> | 56.6 |
| BB12B | Beaver Brook | Sediment | stream | High | <NA> | 56.6 |
| BB13A | Beaver Brook | Sediment | stream | High | <NA> | 56.6 |
| BB14B.raw | Beaver Brook | Sediment | stream | High | <NA> | 56.6 |
| BB21A | Beaver Brook | Sediment | stream | High | <NA> | 48.1 |
| BB21B.raw | Beaver Brook | Sediment | stream | High | <NA> | 48.1 |
| BB22A.raw | Beaver Brook | Water | stream | High | <NA> | 48.1 |
| BB22B | Beaver Brook | Sediment | stream | High | <NA> | 48.1 |
| BB23B.raw | Beaver Brook | Sediment | stream | High | <NA> | 48.1 |
| BB31A.raw | Beaver Brook | Water | stream | High | <NA> | 52 |
| BB31B | Beaver Brook | Sediment | stream | High | <NA> | 52 |
| BB32B | Beaver Brook | Sediment | stream | High | <NA> | 52 |
| BB33A | Beaver Brook | Sediment | stream | High | <NA> | 52 |
| BB34A | Beaver Brook | Sediment | stream | High | <NA> | 52 |
| Coes002 | Coe's Pond | Fecal | pond | Medium | *Lithobates catesbeianus* | 24.7 |
| Coes003 | Coe's Pond | Fecal | pond | Medium | *Lithobates catesbeianus* | 24.7 |
| Coes004 | Coe's Pond | Fecal | pond | Medium | *Lithobates catesbeianus* | 24.7 |
| Coes006 | Coe's Pond | Fecal | pond | Medium | *Lithobates catesbeianus* | 24.7 |
| Coes007 | Coe's Pond | Fecal | pond | Medium | *Lithobates catesbeianus* | 24.7 |
| Coes009 | Coe's Pond | Fecal | pond | Medium | *Lithobates catesbeianus* | 24.7 |
| Coes010 | Coe's Pond | Fecal | pond | Medium | *Lithobates catesbeianus* | 24.7 |
| Cook005 | Cook Pond | Fecal | pond | Medium | *Lithobates catesbeianus* | 17.2 |
| Cook007 | Cook Pond | Fecal | pond | Medium | *Lithobates catesbeianus* | 17.2 |
| Cook008 | Cook Pond | Fecal | pond | Medium | *Lithobates catesbeianus* | 17.2 |
| Cook009 | Cook Pond | Fecal | pond | Medium | *Lithobates catesbeianus* | 17.2 |
| Cook011 | Cook Pond | Fecal | pond | Medium | *Lithobates catesbeianus* | 17.2 |
| Cook012 | Cook Pond | Fecal | pond | Medium | *Lithobates catesbeianus* | 17.2 |
| Cook013 | Cook Pond | Fecal | pond | Medium | *Lithobates catesbeianus* | 17.2 |
| CP21B | Cook Pond | Sediment | stream | Low | <NA> | 9 |
| CP22B | Cook Pond | Sediment | stream | Low | <NA> | 9 |
| CP22J.raw | Cook Pond | Water | stream | Low | <NA> | 9 |
| CP23B | Cook Pond | Sediment | stream | Low | <NA> | 9 |
| CP24A.raw | Cook Pond | Sediment | stream | Low | <NA> | 9 |
| CP31B | Cook Pond | Sediment | stream | Medium | <NA> | 17.2 |
| CP32A.raw | Cook Pond | Sediment | stream | Medium | <NA> | 17.2 |
| CP32J.raw | Cook Pond | Water | stream | Medium | <NA> | 17.2 |
| CP33B | Cook Pond | Sediment | stream | Medium | <NA> | 17.2 |
| CP42J.raw | Cook Pond | Water | pond | Medium | <NA> | 17.2 |
| CR12B | Coe's Pond | Sediment | stream | Medium | <NA> | 24.7 |
| CR21A.raw | Coe's Pond | Water | pond | Medium | <NA> | 24.7 |
| CR31A.raw | Coe's Pond | Water | pond | Medium | <NA> | 24.7 |
| CR32J.raw | Coe's Pond | Water | pond | Medium | <NA> | 24.7 |
| Gl13J.raw | Kinney woods | Water | pond | Low | <NA> | 5.1 |
| KP001 | Kinney woods | Fecal | stream | Low | *Lithobates clamitans* | 5.1 |
| KP002 | Kinney woods | Fecal | stream | Low | *Lithobates clamitans* | 5.1 |
| KP003 | Kinney woods | Fecal | stream | Low | *Lithobates clamitans* | 5.1 |
| KP004 | Kinney woods | Fecal | pond | Low | *Lithobates palustris* | 5.1 |
| KP005 | Kinney woods | Fecal | pond | Low | *Lithobates catesbeianus* | 5.1 |
| KP006 | Kinney woods | Fecal | pond | Low | *Lithobates catesbeianus* | 5.1 |
| KP007 | Kinney woods | Fecal | pond | Low | *Lithobates catesbeianus* | 5.1 |
| KP009 | Kinney woods | Fecal | pond | Low | *Lithobates palustris* | 5.1 |
| KP010 | Kinney woods | Fecal | pond | Low | *Lithobates palustris* | 5.1 |
| KP011 | Kinney woods | Fecal | pond | Low | *Lithobates palustris* | 5.1 |
| KP013 | Kinney woods | Fecal | pond | Low | *Lithobates palustris* | 5.1 |
| MB11B | Holy Cross | Sediment | stream | High | <NA> | 56.7 |
| MB11J.raw | Holy Cross | Water | stream | High | <NA> | 56.7 |
| MB12A.raw | Holy Cross | Sediment | stream | High | <NA> | 56.7 |
| MB13B | Holy Cross | Sediment | stream | High | <NA> | 56.7 |
| MB14A.raw | Holy Cross | Sediment | stream | High | <NA> | 56.7 |
| MP001 | Mill Pond | Fecal | pond | Medium | *Lithobates catesbeianus* | 25.6 |
| MP002 | Mill Pond | Fecal | pond | Medium | *Lithobates clamitans* | 25.6 |
| MP003 | Mill Pond | Fecal | pond | Medium | *Lithobates catesbeianus* | 25.6 |
| MP004 | Mill Pond | Fecal | pond | Medium | *Lithobates catesbeianus* | 25.6 |
| MP005 | Mill Pond | Fecal | pond | Medium | *Lithobates clamitans* | 25.6 |
| MP007 | Mill Pond | Fecal | pond | Medium | *Lithobates clamitans* | 25.6 |
| MP008 | Mill Pond | Fecal | pond | Medium | *Lithobates catesbeianus* | 25.6 |
| MR001 | Middle river | Fecal | stream | High | *Lithobates catesbeianus* | 51.8 |
| MR002 | Middle river | Fecal | stream | High | *Lithobates clamitans* | 51.8 |
| MR003.raw | Middle river | Fecal | stream | High | *Lithobates clamitans* | 51.8 |
| MR004.raw | Middle river | Fecal | stream | High | *Lithobates clamitans* | 51.8 |
| MR005.raw | Middle river | Fecal | stream | High | *Lithobates clamitans* | 51.8 |
| MR007.raw | Middle river | Fecal | stream | High | *Lithobates clamitans* | 51.8 |
| MR010.raw | Middle river | Fecal | stream | High | *Lithobates clamitans* | 51.8 |
| MR011.raw | Middle river | Fecal | stream | High | *Lithobates clamitans* | 51.8 |
| MR012.raw | Middle river | Fecal | stream | High | *Lithobates clamitans* | 51.8 |
| MR014.raw | Middle river | Fecal | stream | High | *Lithobates clamitans* | 51.8 |
| MR015.raw | Middle river | Fecal | stream | High | *Lithobates clamitans* | 51.8 |
| MR11B | Cemetery | Sediment | stream | High | <NA> | 46.2 |
| MR11B.raw | Cemetery | Sediment | stream | High | <NA> | 46.2 |
| MR12B | Cemetery | Sediment | stream | High | <NA> | 46.2 |
| MR12J.raw | Cemetery | Water | stream | High | <NA> | 46.2 |
| MR13A | Cemetery | Sediment | stream | High | <NA> | 46.2 |
| MR14B | Cemetery | Water | stream | High | <NA> | 46.2 |
| MR23A.raw | Middle river | Water | stream | High | <NA> | 51.8 |
| PR001 | Patch reservoir | Fecal | pond | Medium | *Lithobates catesbeianus* | 22 |
| PR002 | Patch reservoir | Fecal | pond | Medium | *Lithobates catesbeianus* | 22 |
| PR003 | Patch reservoir | Fecal | pond | Medium | *Lithobates clamitans* | 22 |
| PR005 | Patch reservoir | Fecal | pond | Medium | *Lithobates catesbeianus* | 22 |
| PR006 | Patch reservoir | Fecal | pond | Medium | *Lithobates clamitans* | 22 |
| PR007 | Patch reservoir | Fecal | pond | Medium | *Lithobates catesbeianus* | 22 |
| PR008 | Patch reservoir | Fecal | pond | Medium | *Lithobates clamitans* | 22 |
| PR009 | Patch reservoir | Fecal | pond | Medium | *Lithobates catesbeianus* | 22 |
| PR010 | Patch reservoir | Fecal | pond | Medium | *Lithobates catesbeianus* | 22 |
| PR11B | Patch reservoir | Sediment | stream | Medium | <NA> | 20.8 |
| PR11J.raw | Patch reservoir | Water | stream | Medium | <NA> | 20.8 |
| PR12B | Patch reservoir | Sediment | stream | Medium | <NA> | 20.8 |
| PR13A | Patch reservoir | Sediment | stream | Medium | <NA> | 20.8 |
| PR14A | Patch reservoir | Sediment | stream | Medium | <NA> | 20.8 |
| PR21J.raw | Patch reservoir | Water | stream | Medium | <NA> | 25.3 |
| PR23B | Patch reservoir | Sediment | stream | Medium | <NA> | 25.3 |
| PR24B | Patch reservoir | Sediment | stream | Medium | <NA> | 25.3 |
| PR33A.raw | Patch reservoir | Water | pond | Medium | <NA> | 22 |
| PR33J.raw | Patch reservoir | Water | pond | Medium | <NA> | 22 |
| PRS001 | Tatnuck Brook | Fecal | stream | Low | *Lithobates catesbeianus* | 9 |
| PRS002 | Tatnuck Brook | Fecal | stream | Low | *Lithobates clamitans* | 9 |
| PRS003 | Tatnuck Brook | Fecal | stream | Low | *Lithobates clamitans* | 9 |
| PRS004 | Tatnuck Brook | Fecal | stream | Low | *Lithobates clamitans* | 9 |
| PRS006 | Tatnuck Brook | Fecal | stream | Low | *Lithobates clamitans* | 9 |
| PRS007 | Tatnuck Brook | Fecal | stream | Low | *Lithobates clamitans* | 9 |
| PRS008 | Tatnuck Brook | Fecal | stream | Low | *Lithobates clamitans* | 9 |
| PRS009 | Tatnuck Brook | Fecal | stream | Low | *Lithobates catesbeianus* | 9 |
| PRS010 | Tatnuck Brook | Fecal | stream | Low | *Lithobates palustris* | 9 |

**Supplementary Table 2**. Relative abundances of the top 20 fungal genera across habitats and urbanization levels.

| **OTU** | **Genus** | **Urbanization Category** | **Is it present?** | **Relative abundance per habitat** | | |
| --- | --- | --- | --- | --- | --- | --- |
|  |  |  |  | **Sediment** | **Water** | **Fecal** |
| ASV10 | *Lemonniera* | High | TRUE | 0.716 | NA | NA |
|  |  | Low | TRUE | 0.295 | NA | NA |
|  |  | Medium | TRUE | 0.306 | NA | NA |
| ASV105 | *Podospora* | High | TRUE | 0.058 | NA | NA |
|  |  | Low | TRUE | 0.252 | NA | NA |
|  |  | Medium | TRUE | 0.018 | NA | NA |
| ASV1140 | *Betamyces* | High | TRUE | NA | 0.083 | NA |
|  |  | Low | FALSE | NA | NA | NA |
|  |  | Medium | TRUE | NA | 0.02 | NA |
| ASV117 | *Hyphoderma* | High | TRUE | 0.008 | NA | NA |
|  |  | Low | FALSE | NA | NA | NA |
|  |  | Medium | TRUE | 0.282 | NA | NA |
| ASV133 | *Lemonniera* | High | TRUE | 0.069 | NA | NA |
|  |  | Low | TRUE | 0.002 | NA | NA |
|  |  | Medium | TRUE | 0.187 | NA | NA |
| ASV140 | *Atractospora* | High | TRUE | 0.009 | NA | NA |
|  |  | Low | TRUE | 0.003 | NA | NA |
|  |  | Medium | TRUE | 0.31 | NA | NA |
| ASV152 | *Hamatocanthoscypha* | High | FALSE | NA | NA | NA |
|  |  | Low | TRUE | 0.231 | NA | NA |
|  |  | Medium | TRUE | 0.021 | NA | NA |
| ASV155 | *Glaciozyma* | High | TRUE | 0.03 | NA | NA |
|  |  | Low | FALSE | NA | NA | NA |
|  |  | Medium | TRUE | 0.295 | NA | NA |
| ASV16 | *Parasarocladium* | High | TRUE | NA | NA | 0.903 |
|  |  | Low | TRUE | NA | NA | 0.019 |
|  |  | Medium | TRUE | NA | NA | 0.909 |
| ASV170 | *Coleophoma* | High | TRUE | NA | 0.008 | NA |
|  |  | Low | TRUE | NA | 0.137 | NA |
|  |  | Medium | TRUE | NA | 0.203 | NA |
| ASV18 | *Stagonosporopsis* | High | TRUE | NA | NA | 0.01 |
|  |  | Low | TRUE | NA | NA | 0.579 |
|  |  | Medium | TRUE | NA | NA | 0.112 |
| ASV195 | *Basidiobolus* | High | FALSE | NA | NA | NA |
|  |  | Low | FALSE | NA | NA | NA |
|  |  | Medium | TRUE | NA | 0.143 | NA |
| ASV2 | *Cladosporium* | High | TRUE | NA | NA | 0.607 |
|  |  | Low | TRUE | NA | NA | 0.244 |
|  |  | Medium | TRUE | NA | NA | 1.701 |
| ASV20 | *Alternaria* | High | TRUE | NA | NA | 0.211 |
|  |  | Low | TRUE | NA | NA | 0.011 |
|  |  | Medium | TRUE | NA | NA | 0.382 |
| ASV21 | *Plectosphaerella* | High | TRUE | NA | NA | 0.808 |
|  |  | Low | TRUE | NA | NA | 0.028 |
|  |  | Medium | TRUE | NA | NA | 0.358 |
| ASV215 | *Candida* | High | TRUE | NA | 0.17 | NA |
|  |  | Low | TRUE | NA | 0.015 | NA |
|  |  | Medium | TRUE | NA | 0.054 | NA |
| ASV24 | *Didymella* | High | TRUE | NA | NA | 0.041 |
|  |  | Low | TRUE | NA | NA | 0.577 |
|  |  | Medium | TRUE | NA | NA | 0.344 |
| ASV26 | *Plectosphaerella* | High | TRUE | NA | NA | 0.056 |
|  |  | Low | TRUE | NA | NA | 0.6 |
|  |  | Medium | TRUE | NA | NA | 0.211 |
| ASV28 | *Basidiobolus* | High | TRUE | NA | 0.055 | NA |
|  |  | Low | TRUE | NA | 0.04 | NA |
|  |  | Medium | TRUE | NA | 0.42 | NA |
| ASV29 | *Aureobasidium* | High | TRUE | NA | 0.791 | 0.082 |
|  |  | Low | TRUE | NA | 0.008 | 0.158 |
|  |  | Medium | TRUE | NA | 0.176 | 0.417 |
| ASV291 | *Basidiobolus* | High | FALSE | NA | NA | NA |
|  |  | Low | FALSE | NA | NA | NA |
|  |  | Medium | TRUE | NA | 0.088 | NA |
| ASV3 | *Basidiobolus* | High | TRUE | NA | NA | 0.016 |
|  |  | Low | TRUE | NA | NA | 0.849 |
|  |  | Medium | TRUE | NA | NA | 0.178 |
| ASV31 | *Pyrenochaetopsis* | High | TRUE | 0.23 | NA | NA |
|  |  | Low | TRUE | 0.005 | NA | NA |
|  |  | Medium | TRUE | 0.12 | NA | NA |
| ASV35 | *Anguillospora* | High | TRUE | 0.769 | 0.203 | NA |
|  |  | Low | TRUE | 0.095 | 0.01 | NA |
|  |  | Medium | TRUE | 0.311 | 0.105 | NA |
| ASV37 | *Phaeosphaeria* | High | TRUE | 0.337 | NA | NA |
|  |  | Low | TRUE | 0.004 | NA | NA |
|  |  | Medium | TRUE | 0.063 | NA | NA |
| ASV38 | *Botrytis* | High | TRUE | NA | NA | 0.836 |
|  |  | Low | TRUE | NA | NA | 0.001 |
|  |  | Medium | TRUE | NA | NA | 0.007 |
| ASV39 | *Basidiobolus* | High | TRUE | NA | NA | 0.001 |
|  |  | Low | TRUE | NA | NA | 0.537 |
|  |  | Medium | TRUE | NA | NA | 0.052 |
| ASV4 | *Basidiobolus* | High | TRUE | NA | NA | 0.003 |
|  |  | Low | TRUE | NA | NA | 1.635 |
|  |  | Medium | TRUE | NA | NA | 0.252 |
| ASV41 | *Candida* | High | TRUE | 0.735 | 0.208 | NA |
|  |  | Low | TRUE | 0.022 | 0.09 | NA |
|  |  | Medium | TRUE | 0.043 | 0.341 | NA |
| ASV43 | *Mrakia* | High | TRUE | 0.115 | NA | NA |
|  |  | Low | TRUE | 0.003 | NA | NA |
|  |  | Medium | TRUE | 0.29 | NA | NA |
| ASV44 | *Chloridium* | High | FALSE | NA | NA | NA |
|  |  | Low | TRUE | NA | NA | 0.517 |
|  |  | Medium | TRUE | NA | NA | 0.179 |
| ASV48 | *Cladosporium* | High | TRUE | NA | NA | 0.544 |
|  |  | Low | TRUE | NA | NA | 0.002 |
|  |  | Medium | TRUE | NA | NA | 0.193 |
| ASV489 | *Zygophlyctis* | High | FALSE | NA | NA | NA |
|  |  | Low | TRUE | NA | 0.182 | NA |
|  |  | Medium | TRUE | NA | 0.344 | NA |
| ASV5 | *Basidiobolus* | High | TRUE | NA | NA | 0.031 |
|  |  | Low | TRUE | NA | NA | 1.57 |
|  |  | Medium | TRUE | NA | NA | 0.472 |
| ASV509 | *Aquanectria* | High | TRUE | NA | 0.255 | NA |
|  |  | Low | TRUE | NA | 0.003 | NA |
|  |  | Medium | TRUE | NA | 0.022 | NA |
| ASV52 | *Fusarium* | High | TRUE | NA | NA | 0.001 |
|  |  | Low | FALSE | NA | NA | NA |
|  |  | Medium | TRUE | NA | NA | 0.617 |
| ASV525 | *Basidiobolus* | High | FALSE | NA | NA | NA |
|  |  | Low | TRUE | NA | 0.394 | NA |
|  |  | Medium | TRUE | NA | 0.024 | NA |
| ASV54 | *Solicoccozyma* | High | TRUE | 0.116 | NA | NA |
|  |  | Low | TRUE | 0.025 | NA | NA |
|  |  | Medium | TRUE | 0.54 | NA | NA |
| ASV55 | *Saitozyma* | High | TRUE | 0.131 | NA | NA |
|  |  | Low | TRUE | 0.116 | NA | NA |
|  |  | Medium | TRUE | 0.11 | NA | NA |
| ASV56 | *Clavispora* | High | TRUE | NA | 0.167 | NA |
|  |  | Low | TRUE | NA | 0.219 | NA |
|  |  | Medium | TRUE | NA | 1.062 | NA |
| ASV58 | *Glaciozyma* | High | TRUE | NA | NA | 0.005 |
|  |  | Low | TRUE | NA | NA | 0.033 |
|  |  | Medium | TRUE | NA | NA | 0.962 |
| ASV59 | *Conocybe* | High | TRUE | 0.513 | NA | NA |
|  |  | Low | FALSE | NA | NA | NA |
|  |  | Medium | TRUE | 0.22 | NA | NA |
| ASV6 | *Basidiobolus* | High | TRUE | NA | NA | 0.004 |
|  |  | Low | TRUE | NA | NA | 1.529 |
|  |  | Medium | TRUE | NA | NA | 0.272 |
| ASV610 | *Rhytisma* | High | TRUE | NA | 0.171 | NA |
|  |  | Low | FALSE | NA | NA | NA |
|  |  | Medium | TRUE | NA | 0.029 | NA |
| ASV66 | *Metarhizium* | High | TRUE | NA | 0.62 | NA |
|  |  | Low | FALSE | NA | NA | NA |
|  |  | Medium | TRUE | NA | 0.089 | NA |
| ASV663 | *Betamyces* | High | TRUE | NA | 0.062 | NA |
|  |  | Low | TRUE | NA | 0.038 | NA |
|  |  | Medium | TRUE | NA | 0.155 | NA |
| ASV7 | *Candida* | High | TRUE | NA | NA | 0.057 |
|  |  | Low | TRUE | NA | NA | 1.01 |
|  |  | Medium | TRUE | NA | NA | 0.085 |
| ASV70 | *Tausonia* | High | TRUE | 0.449 | NA | NA |
|  |  | Low | TRUE | 0.003 | NA | NA |
|  |  | Medium | TRUE | 0.066 | NA | NA |
| ASV725 | *Rhizophydium* | High | FALSE | NA | NA | NA |
|  |  | Low | FALSE | NA | NA | NA |
|  |  | Medium | TRUE | NA | 0.134 | NA |
| ASV74 | *Dichostereum* | High | FALSE | NA | NA | NA |
|  |  | Low | TRUE | NA | NA | 0.366 |
|  |  | Medium | TRUE | NA | NA | 0.031 |
| ASV75 | *Apodus* | High | TRUE | 0.078 | NA | NA |
|  |  | Low | TRUE | 0.319 | NA | NA |
|  |  | Medium | TRUE | 0.036 | NA | NA |
| ASV76 | *Tetrachaetum* | High | TRUE | 0.541 | NA | NA |
|  |  | Low | TRUE | 0.018 | NA | NA |
|  |  | Medium | TRUE | 0.143 | NA | NA |
| ASV78 | *Basidiobolus* | High | FALSE | NA | NA | NA |
|  |  | Low | TRUE | NA | 0.025 | NA |
|  |  | Medium | TRUE | NA | 0.462 | NA |
| ASV845 | *Kazachstania* | High | TRUE | NA | 0.115 | NA |
|  |  | Low | FALSE | NA | NA | NA |
|  |  | Medium | TRUE | NA | 0.012 | NA |
| ASV862 | *Betamyces* | High | TRUE | NA | 0.021 | NA |
|  |  | Low | FALSE | NA | NA | NA |
|  |  | Medium | TRUE | NA | 0.145 | NA |
| ASV9 | *Neoascochyta* | High | TRUE | 0.994 | NA | NA |
|  |  | Low | TRUE | 0.01 | NA | NA |
|  |  | Medium | TRUE | 0.119 | NA | NA |
| ASV94 | *Chaetomella* | High | TRUE | 0.069 | NA | NA |
|  |  | Low | TRUE | 0.007 | NA | NA |
|  |  | Medium | TRUE | 0.355 | NA | NA |

**Supplementary Table 3**. Kruskal-Wallis test statistics for fungal trophic guild abundances across urbanization levels in each habitat.

| **Habitat** | **Guild** | **Chi-square** | **DF** | **p value** |
| --- | --- | --- | --- | --- |
| **Fecal** | Pathogen | 2.9 | 2 | 0.24 |
|  | Symbiont | 2.4 | 2 | 0.23 |
|  | Saprotroph | 2.3 | 2 | 0.33 |
| **Sediment** | Pathogen | 4.0 | 2 | 0.14 |
|  | Symbiont | 3.1 | 2 | 0.21 |
|  | Saprotroph | 1.1 | 2 | 0.57 |
| **Water** | Pathogen | 0.6 | 2 | 0.74 |
|  | Symbiont | 1.8 | 2 | 0.41 |
|  | Saprotroph | 5.8 | 2 | 0.06 |

**Supplementary Table 4**. Summary of permutation ANOVA and pairwise permutation test results for fungal trophic guild abundances across urbanization levels in fecal, sediment, and water habitats.

| **Habitat** | **Guild** | **DF** | **Sum Sq** | **Mean Sq** | **F value** | **Pr(>F)** | **Pairwise permutations** | | |
| --- | --- | --- | --- | --- | --- | --- | --- | --- | --- |
|  |  |  |  |  |  |  | **High - Low (p adjusted)** | **High - Medium (p adjusted)** | **Low - Medium (p adjusted)** |
| Fecal | Pathotroph | 2 | 2685731082 | 1342865541 | 4.276 | 0.102 | 0.365 | 0.365 | 0.2155 |
|  | Symbiotroph | 2 | 457117540 | 228558770 | 2.026 | 0.247 | 0.299 | 0.299 | 0.299 |
|  | Saprotroph | 2 | 868926320 | 434463160 | 0.309 | 0.750 | 0.926 | 0.926 | 0.926 |
| Sediment | Pathotroph | 2 | 2.12E-03 | 1.06E-03 | 2.419 | 0.205 | 0.254 | 0.254 | 0.5318 |
|  | Symbiotroph | 2 | 6.80E-05 | 3.40E-05 | 6.804 | 0.052 | 0.193 | 0.162 | 0.864 |
|  | Saprotroph | 2 | 7.58E-03 | 3.79E-03 | 1.189 | 0.393 | 0.906 | 0.431 | 0.431 |
| Water | Pathotroph | 2 | 0.00022852 | 0.00011426 | 0.091 | 0.914 | 0.938 | 0.938 | 0.938 |
|  | Symbiotroph | 2 | 0.00021613 | 0.00010806 | 0.509 | 0.625 | 0.574 | 0.574 | 0.574 |
|  | Saprotroph | 2 | 0.02485638 | 0.01242819 | 6.115 | 0.036 | 0.243 | 0.163 | 0.163 |

**Supplementary files**

**Supplementary File 1**. Differential abundant ASV for fecal samples using DESeq2 comparing ASV abundance at low urbanization versus high urbanization. The table includes the following columns: baseMean, representing the average normalized expression value across all samples; log2FoldChange, where positive values indicate significant abundance at low urbanization, and negative values values indicate significant abundance at high urbanization; lfcSE, the standard error of the log₂ fold change estimate; stat, the Wald statistic for differential expression testing; pvalue, the raw p-value for the significance of the observed expression difference; and padj, the adjusted p-value (using the Benjamini-Hochberg method) to control for the false discovery rate.

**Supplementary File 2**. Differential abundant ASV for sediment samples using DESeq2 comparing ASV abundance at low urbanization versus high urbanization. The table includes the following columns: baseMean, representing the average normalized expression value across all samples; log2FoldChange, where positive values indicate significant abundance at low urbanization, and negative values values indicate significant abundance at high urbanization; lfcSE, the standard error of the log₂ fold change estimate; stat, the Wald statistic for differential expression testing; pvalue, the raw p-value for the significance of the observed expression difference; and padj, the adjusted p-value (using the Benjamini-Hochberg method) to control for the false discovery rate.

**Supplementary File 3**. Differential abundant ASV for water samples using DESeq2 comparing planktonic ASV abundance at low urbanization versus high urbanization. The table includes the following columns: baseMean, representing the average normalized expression value across all samples; log2FoldChange, where positive values indicate significant abundance at low urbanization, and negative values values indicate significant abundance at high urbanization; lfcSE, the standard error of the log₂ fold change estimate; stat, the Wald statistic for differential expression testing; pvalue, the raw p-value for the significance of the observed expression difference; and padj, the adjusted p-value (using the Benjamini-Hochberg method) to control for the false discovery rate.
